# MDQ: A DockQ-Inspired Scoring Function for Assessing Interface Similarity and Clustering in Molecular Dynamics

**DOI:** 10.64898/2026.11.19.767893

**Authors:** Jenny Ehles, Florent Barbault, Tom Miclot

## Abstract

Assessing the similarity between protein–protein interfaces is essential for evaluating docking models, yet widely used metrics such as DockQ rely on a single native reference structure and yield asymmetric scores, making them unsuitable for molecular dynamics (MD) trajectories where no reference exists and frame-to-frame comparisons are required. Here, we introduce the MDQ, a fully symmetric reformulation of DockQ designed for MD trajectory analysis. MDQ replaces the fraction of native contacts with the Fraction of Shared Contacts (FSC), and redefines interface and ligand RMSD as weighted variants that penalize low atomic overlap between compared frames. The score can be used to follow structure/contacts convergence of an interface along a trajectory. And it is inverted to produce a true distance metric ranging from 0 (identical interfaces) to 1 (distinct interfaces), directly compatible with distance-based clustering. Applied to two benchmark complexes, Ail-Vitronectin and TGFBR1-FKBP12, MDQ heatmaps reveal global interface stability alongside subtle conformational rearrangements, while DBSCAN and hierarchical clustering of the MDQ distance matrix identifies representative conformational states. Then non-bonding interaction found in clusters were identified by MICLOT software. MDQ thus provides a robust and frame-to-frame symmetric framework for monitoring protein-protein interface conformational and interaction heterogeneity throughout MD simulations.

**VISUAL ABSTRACT:** 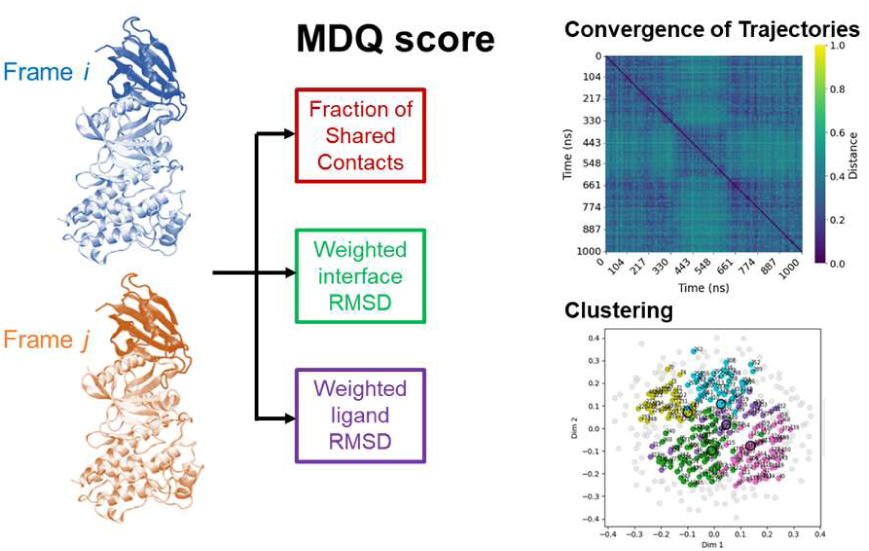

## 1. INTRODUCTION

Protein-protein interactions (PPIs) are fundamental to numerous cellular processes, relying on geometric and physicochemical complementarity between binding partners. Understanding their association mechanisms is not only critical for understanding biological function but also holds significant therapeutic potential, as PPIs represent key targets for drug design. Molecular dynamics (MD) simulations, combined with experimental validation, provide a powerful approach to investigate the formation, stability, and conformational dynamics of protein interfaces.

Traditionally, the structural dynamics of proteins or the clustering of conformations have been analyzed using the Root Mean Square Deviation (RMSD), typically computed on Cα atoms of the backbone or all heavy atoms. Over the years, other metrics have emerged to assess structural similarity, including MaxSub^1^, TM-score^2^, Global Distance Test^3^, S-score^4^, IS-score^5^, Local-Global Alignment^6^, Global Distance Calculation^7^, many of which are widely used in docking to evaluate predicted models against experimental structures.

Among these, DockQ^8,9^ has become one of the most widely adopted metrics in docking, including for AI-predicted protein complexes. Based on the CAPRI^10–12^ (Critical Assessment of PRedicted Interactions) criteria—such as the fraction of native and non-native contacts, ligand RMSD (LRMSD), and interface RMSD (IRMSD). DockQ integrates these descriptors into a single score ranging from 0, for poor model, to 1, forhigh-quality model. This bounded range enables classification according to CAPRI categories^8,9,13^: Incorrect (0.00 ≤ DockQ < 0.23), Acceptable (0.23 ≤ DockQ < 0.49), Medium (0.49 ≤ DockQ < 0.80), and High quality (DockQ ≥ 0.80). Its scalability has also made it suitable for machine learning applications, such as pDockQ^14,15^. Furthermore, integrating CAPRI-defined parameters into MD simulations allows for the refinement and evaluation of predicted models^16,17^, while tools like DynaPIN^18^ integrate DockQ to track interface dynamics relative to a reference structure.

However, despite its utility, DockQ has a critical limitation in MD trajectory analysis: it is inherently asymmetric, as it is designed to compare a model against a reference structure, answering “how similar” the model is to the reference. Consequently, comparing structure *i* to *j* does not yield the same score as comparing *j* to *i*, unlike mathematically symmetric metrics such as RMSD. More explanations are in SI, sections I. This asymmetry restricts its applicability for analysing the dynamic evolution of interfaces over time.

To address this limitation, we introduce MDQ, a symmetric redefinition of DockQ that retains its original formulation but ensures consistency regardless of the order of comparison (see Figure 1). Obviously, MDQ enables the tracking of interface dynamics relative to a reference, providing a score variation curve across trajectory frames. Due to its symmetric nature, MDQ also allows the generation of 2D maps analogous to 2D-RMSD, facilitating the monitoring of interface convergence or divergence along a trajectory. Additionally, MDQ’s inverted scoring scheme where 0 indicates similar interfaces and 1 indicates dissimilar interfaces, to makes it compatible with clustering algorithms. The resulting score matrix can thus be used not only for dynamic mapping but also to identify distinct interface conformations present in the trajectory.

**Figure 1.**
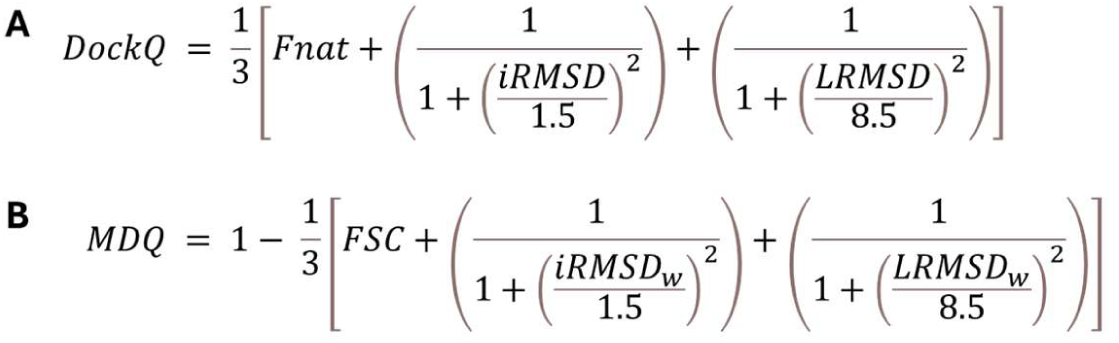
Comparison between the original DockQ equation (A) and the proposed symmetric MDQ formulation (B). Panel A shows the standard DockQ score combining the Fnat, interface iRMSD, and LRMSD, with scaling parameters d₁ = 1.5 Å and d₂ = 8.5 Å. Panel B presents MDQ, which replaces Fnat with the symmetric fraction of shared contacts (FSC) and uses weighted RMSD values corrected for atomic overlap. The leading “1 −” is used to inverts the score scale so that MDQ = 0 indicates identical interfaces and MDQ close to 1 indicates completely dissimilar interfaces, reversing the DockQ convention. Both formulations retain the original d₁ and d₂ parameters.

In this work, we apply MDQ to analyse two protein complexes with stable interfaces. The results demonstrate that MDQ effectively captures interface stability through 2D maps and is sufficiently sensitive to reveal fine conformational variations adopted by each interface during the simulations. This sensitivity is further supported by an analysis of non-bonding interactions for each representative structure of the identified clusters, revealing subtle differences in the interaction networks established within the interfaces.

## 2. METHODS

### 2.1. Fraction of shared contact (FSC)

#### 2.1.a. Detection of pair of residues

A pair of residues is considered in contact if at least one pair of their heavy atoms is within a distance of 2 Å < d_i,j_ < 5 Å. The upper threshold of 5 Å follows the CAPRI^10–12^ recommendations for defining biologically relevant contacts. To exclude non-physiological clashes resulting from modelling errors, a lower distance limit of 2 Å is applied, a value commonly used in modelling software^19^. While alternative approaches employ stricter thresholds, up to 0.4 Å for atomic overlaps^20,21^, the 2 Å cutoff was selected to balance biological relevance with computational robustness.

#### 2.1.b. Computing the fraction of shared contact

A list of residue pairs in contact is established for each analysed conformation. From these lists, the common elements are identified, representing the shared contacts between the two conformations (see Figure 2). The FSC is then defined as the ratio of twice the number of common contacts to the total number of contacts identified in both structures. Since common contacts are present in both structures, their count is doubled to reflect their shared nature. This leads to the Dice-Sørensen coefficient^22,23^ equation:

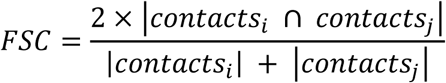

**Figure 2.**
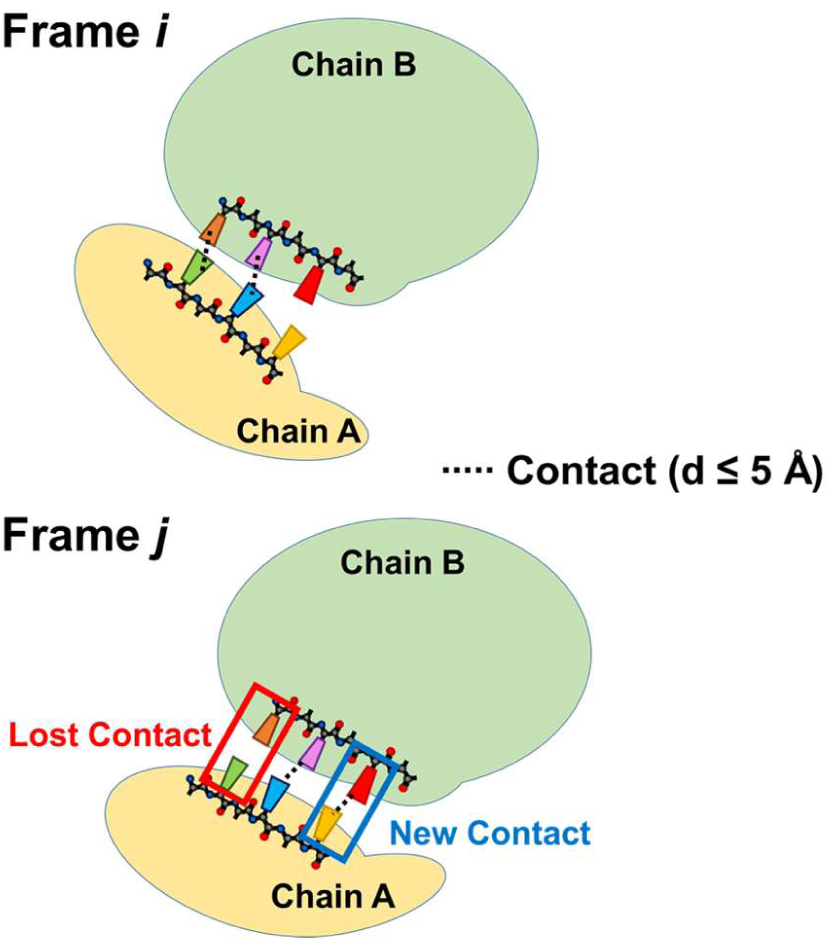
Schematic illustration of inter-chain residue contacts between two protein chains (A and B) in frames *i* and *j* of a MD trajectory. A contact is defined when at least one pair of heavy atoms from the two residues are within 5 Å (dashed lines), following CAPRI recommendations. Frame *j* highlights contacts lost (red box) and gained (blue box) relative to frame *i*, illustrating the dynamic reorganization of the protein-protein interface along the trajectory.

This formulation distinguishes our approach from the Fraction of Native Contacts (fnat) defined by CAPRI^10–12^ and used in DockQ^8,9^ or FCC^24^, which typically compares contacts to a reference native structure rather than between two arbitrary conformations.

### 2.2. Computing RMSD values

#### 2.2.a. Detection ligand and receptor atoms

The iRMSD and LRMSD are calculated using backbone heavy atoms of the proteins. First, atoms within 10 Å of each other between the ligand and receptor are selected to define the interface atoms. To ensure proper structural alignment, only atoms common to both frames (i and j) are retained before RMSD computation.

#### 2.2.a. Computing iRMSD and LRMSD

For iRMSD, both alignment and RMSD calculation are performed on all identified ligand and receptor atoms (see Figure 3). For LRMSD, alignment is restricted to the receptor atoms, while the RMSD is computed exclusively on the ligand atoms (see Figure 3), corresponding to the pocket-aligned LRMSD as defined in DockQ v2^9^. For example, this approach could be particularly suited for 1) peptide interfaces, such as very small peptide studies, 2) localized but non-contiguous interfaces distributed across multiple regions or 3) systems involving flexible or disordered regions undergoing conformational changes.

**Figure 3.**
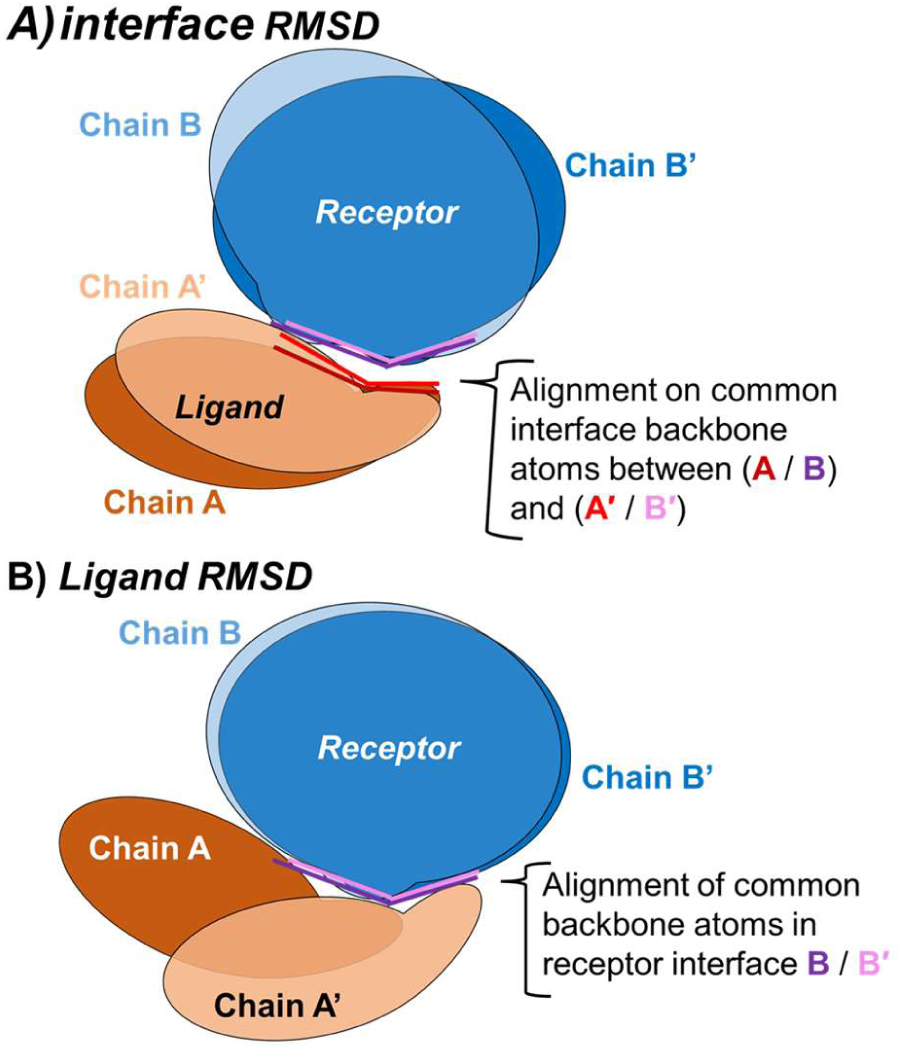
Interface atoms are first selected as those within 10 Å between the ligand and receptor. **(A)** For iRMSD, structural alignment and RMSD calculation are performed on the common interface backbone atoms of both the ligand (A in frame *i* / A′ in frame *j*) and receptor (B in frame *i* / B′ in frame *j*). **(B)** In the case of the LRMSD, alignment is restricted to the common receptor backbone atoms (B/B′), while the RMSD is computed exclusively on the ligand interface atoms (A/A′).

### 2.3. Correction of RMSD values

A similar issue arises with interface atoms as observed with residue contact pairs. Due to the dynamic nature of trajectories, the ligand and receptor atoms forming the interface can vary between frames (see Figure 4). Since alignment and RMSD calculation require identical atoms across different frames, only atoms common to both interface structures are considered. However, this introduces a new problem: a very small RMSD value may result from alignment and calculation over only a few shared atoms, leading to misleading results. To address this, the RMSD values are corrected using a loss function that take in accounts the fraction of shared atoms. The shared atom fraction (frac(i,j)) is calculated in a manner analogous to the FSR. frac(i,j)) is defined as a Dice-Sørensen coefficient, where atoms are the same as those identified before calculating the iRMSD or LRMSD in structures of frame *i* and frame *j*:

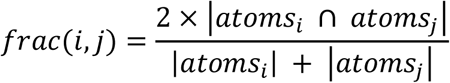

**Figure 4.**
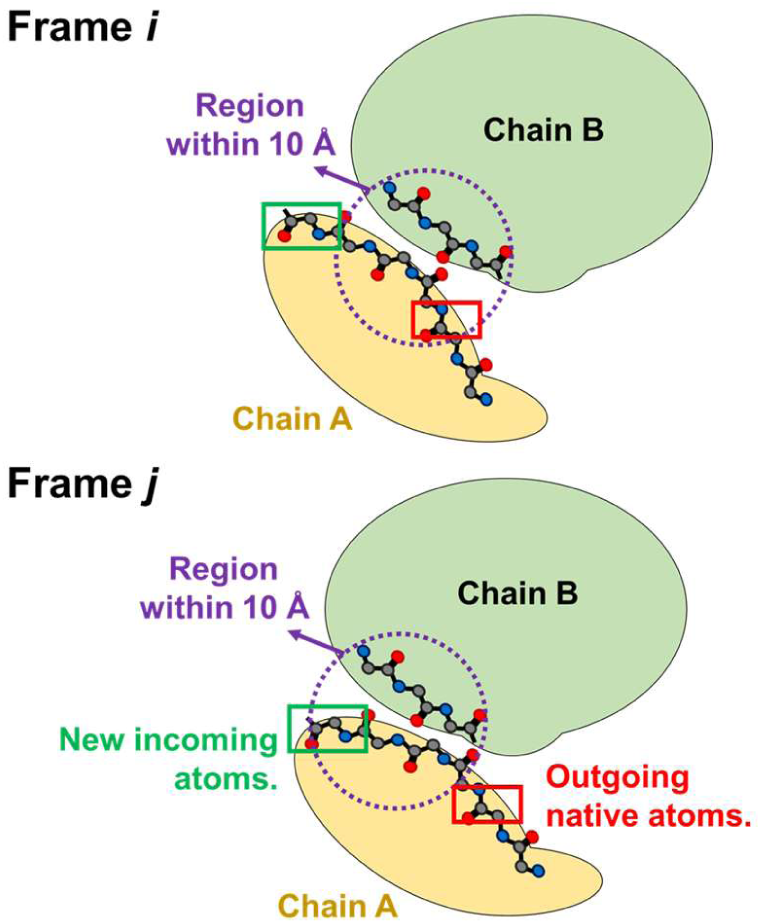
Dynamic variability of interface atoms between two MD trajectory frames (*i* and *j*). The interface region is defined as all atoms within 10 Å of the opposing chain (purple dashed circle). Between frames, the atom composition of the interface changes: new atoms enter the interface (green box) while others leave (red box). Because structural alignment and RMSD calculation require identical atoms across frames, only atoms common to both interface of frame *i* and *j* are retained, motivating the correction of RMSD values based on the fraction of shared atoms.

The frac(i,j) value is then incorporated into the loss function, which follows a logarithmic form. This function approaches 1 as frac(i,j) approaches 1, ensuring minimal correction when all interface atoms are shared. Conversely, it assigns increasingly higher values as frac(i,j) approaches 0, strongly penalizing cases where only a small subset of atoms is shared. Through this approach, the RMSD remains uncorrected when all interface atoms are common, is moderately corrected when most atoms are shared, and is heavily penalized when only a few atoms overlap.

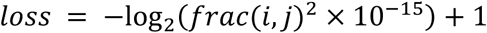

The corrected RMSD (weighted RMSD: RMSD_w_) is computed as the product of the raw RMSD and the loss function:

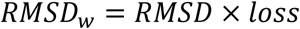

A critical discussion of this method is provided in the SI, sections II and III.

### 2.4. Interpreting MDQ Score results

The MDQ score follows an inverse interpretation compared to DockQ to align with clustering methods, where a score of 0 indicates identical structures and 1 represents maximal dissimilarity. The classification of score values is based on the one used by the DockQ score^8,9^ and globally align with CAPRI assessment^13^. Specifically, a MDQ score greater than 0.80 corresponds to very low similarity (i.e. “Incorrect” in CAPRI), while scores between 0.60 and 0.80 indicate low similarity (i.e. “Acceptable” in CAPRI). A range of 0.40 to 0.60 reflects moderate similarity (i.e. “Medium” in CAPRI), and scores from 0.20 to 0.40 denote high similarity (i.e. “High quality” in CAPRI). Finally, a MDQ score below 0.20 signifies very high similarity between the compared structures (i.e. “High quality” in CAPRI, again).

### 2.5. All-atoms molecular dynamics

#### 2.5.a. Ail/vitronectin complex

Trajectories were previously generated in previously published work^25^. This system was selected due to its relevance as a membrane protein complex, which aligns with the focus of our analysis. The Ail/vitronectin complex has been characterized in this work, demonstrating stability over long-scale molecular dynamics simulations. By leveraging these pre-existing trajectories, we can directly compare our analytical methods with the findings of this study, ensuring consistency and reliability in our assessments. The stability and well-documented behaviour ^25^ of this complex provide a robust foundation for validating our approach.

#### 2.5.b. Cytoplasmic domain of the type I TGF beta receptor / FKBP12complex

The molecular dynamics simulations were based on the crystal structure of the protein solved by X-ray diffraction at a resolution of 2.60 Å, PDB accession: 1B6C^26^, which is part of the CAPRI-Q^13^ benchmark dataset for model validation purposes. The system was prepared using OpenMM-Setup^27^. All heterogens and hydrogens were initially removed from the crystal structure. Missing residues and heavy atoms were reconstructed and added using the integrated tools. Hydrogens were then reintroduced to match a pH of 7.0. The protein was solvated in a cubic water box with a padding distance of 10 Å. Ions were added to neutralize the system, and an ionic strength of 0.5 M NaCl was set. The AMBER14SB^28^ force field was used to describe the protein, while the TIP3P-FB^29^ model was employed for the solvent. To accelerate the simulations, Hydrogen Mass Repartitioning (HMR)^30^ was applied, along with rigid water constraints and all-bond constraints. Non-bonded interactions were treated using the Particle Mesh Ewald (PME)^31^ method with a cutoff distance of 1.0 nm. Temperature was set constant at 300 K. The simulation protocol began with an energy minimization step, followed by a 50 ns NVT equilibration with a 2 fs timestep. This was followed by an additional 50 ns NPT equilibration, also with a 2 fs timestep. Finally, a 1 µs production run was performed in the NPT ensemble with a 4 fs timestep, enabled by HMR and constraints. Three independent replica trajectories were computed using OpenMM^27^ and stored on the Jean Zay supercomputer (V100 nodes) at the French national computing cluster.

### 2.6. Follow the temporal evolution of the system in the MD trajectory

The MDQ score is computed between all frames of a trajectory to assess its temporal evolution. This allows the construction of a 2D heatmap of MDQ scores, visualizing how the structural similarity of a given frame compares to all others. Analogous to 2D-RMSD analysis but with greater precision, this approach enables the detection of conformational changes and/or interface convergence throughout the trajectory. Each replicate is analysed independently of the others.

### 2.7. Interface cluster analysis of the MD trajectory

### 2.7. a. Identify and visualise clusters

The clustering analysis aims to identify distinct interface conformations across the simulated trajectories. Replicates are first merged into a single trajectory, and the MDQ scores are computed for all frame-to-frame comparisons, resulting in a similarity matrix. This matrix is then subjected to clustering algorithms, specifically DBSCAN^32^ and Hierarchical clustering^33^, using the Scikit-learn^34^ implementation, to group structurally similar frames.

#### 2.7.b. Evaluating similarity between identified clusters

For each identified cluster, a representative structure is selected. It corresponds to the frame with the lowest average MDQ score relative to all other frames within the same cluster, ensuring it is the most central and representative conformation. To assess the similarity between clusters, the MDQ scores between their representative structures are evaluated: clusters with low MDQ scores are considered structurally close, while high MDQ scores indicate significant structural divergence.

#### 2.7.b. Visualising the clusters

Given the high-dimensional nature of the data, direct visualization is challenging. To address this, manifold learning techniques: Multi-dimensional Scaling (MDS)^35^, t-distributed Stochastic Neighbor Embedding (t-SNE)^36^, and Uniform Manifold Approximation and Projection (UMAP)^37^ are employed to project the clusters into lower-dimensional spaces. However, the choice of visualization method can significantly influence the perceived and interpreted spatial relationships between clusters.

### 2.8. Analysis of non-bonding interactions

The most representative structures from each cluster, as identified by DBSCAN or Hierarchical, are analysed using MICLOT^38^ software to determine the non-bonding interactions that stabilise the various interfaces.

## 3. RESULTS AND VALIDATION

To validate the sensitivity of MDQ, we deliberately selected stable systems. The rationale is that if the score can detect subtle conformational changes and resolve distinct clusters within systems known to be stable, it demonstrates very good precision. Indeed, both investigated systems, MDQ analysis reveals meaningful differences in interface dynamics and successfully identifies distinct conformational clusters, underscoring the method’s ability to capture fine-scale structural variations that would otherwise remain undetected.

### 3.1. Ail-Vitronectin complex

#### 3.1.a. Stability of the interface

The formation of the Ail-Vitronectin complex has been demonstrated experimentally^39,40^, and an all-atom structural model (see Figure 5A) was subsequently generated using MD simulations^25^. The 2D MDQ heatmap (Figure 5B), computed between all frames of the production phase (i.e., after excluding minimization and equilibration steps), provides a global overview of interface similarity across the merged trajectory. The matrix is symmetric about the diagonal, with identical motifs on either side of it. Intermediate MDQ values (0.4-0.6, green) are observed between the initial and final frames of each replicate, indicating that interfaces sampled in different replicates remain globally comparable. Along the diagonal blocks, MDQ values are predominantly low (0.2-0.4, blue), reflecting high conformational homogeneity and the absence of major structural transitions. Overall, the low MDQ values across the heatmap demonstrate that the Ail-Vitronectin interface is globally stable and structurally converged.

**Figure 5.**
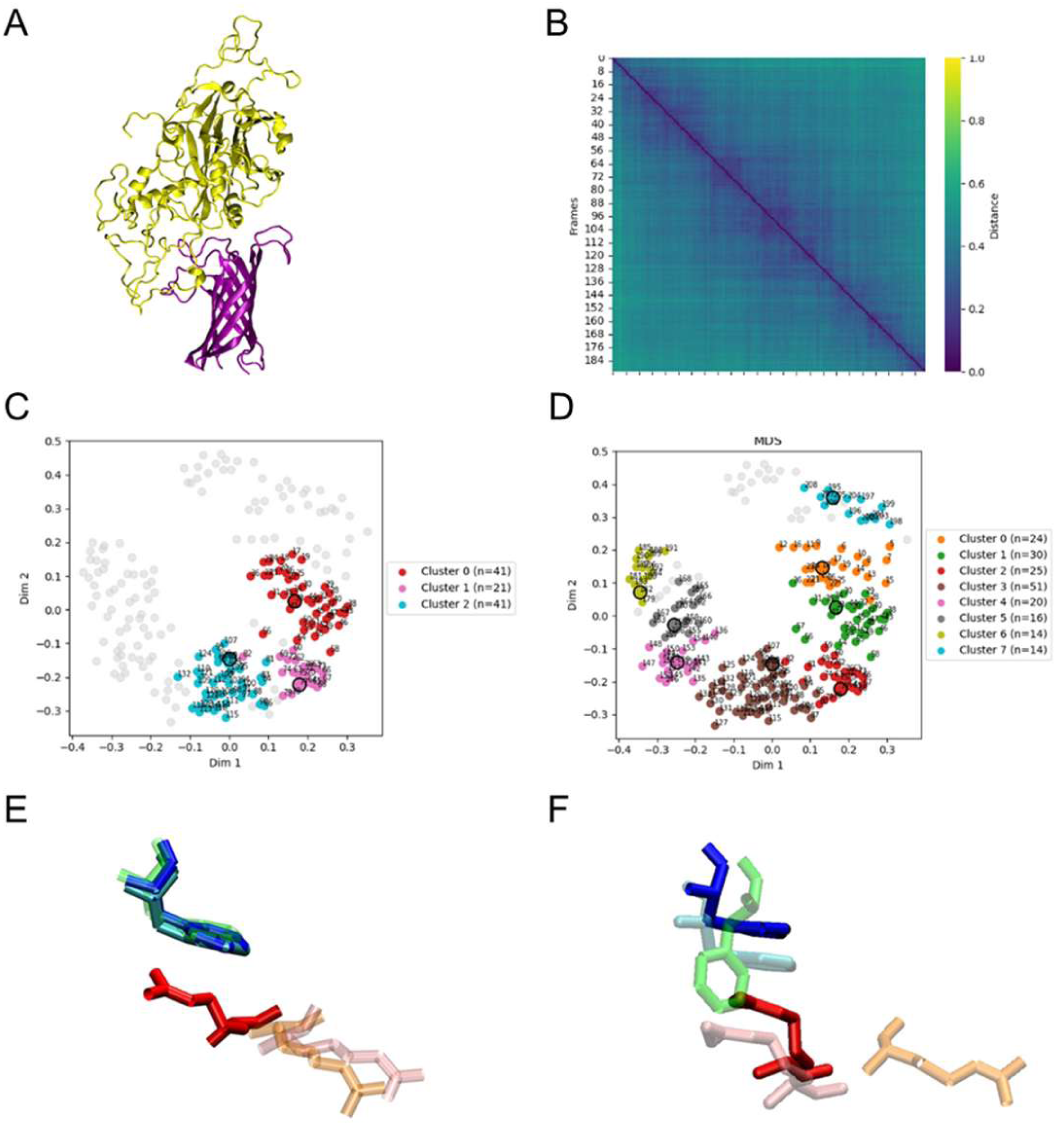
Conformational analysis of the Ail–vitronectin complex. **(A)** 3D structure of the complex (Ail in purple, vitronectin in yellow). **(B)** 2D heatmap of the MDQ score matrix computed between all trajectory frames, where values close to 0 (blue) indicate highly similar interfaces and values close to 1 (yellow) indicate a stability along the trajectory (replica 1). **(C, D)** MDS projections of the trajectory frames colored according to **(C)** DBSCAN clustering (3 clusters) and **(D)** hierarchical clustering (8 clusters), representative structure of each cluster is marked with a black circle. **(E, F)** Close-up views of an Arg-Aromatic interaction: the Arg374-Phe54 pairs found **(E)** in the 3 DBSCAN clusters and **(F)** in selected 3 most different arrangement clusters, for visibility.

#### 3.1.b. Conformational organization and clustering

To resolve the conformational landscape of the Ail–vitronectin interface, we applied two complementary clustering strategies to the symmetric MDQ distance matrix: DBSCAN (see Figure 5C and S5), a density-based method that does not require prior specification of the cluster number, and agglomerative hierarchical clustering (see Figure 5D and S5), which builds a nested partition of the data. DBSCAN resolved the merged trajectory into three clusters, whereas hierarchical clustering partitioned the same frames into eight smaller groups. The clusters identified by DBSCAN were fully contained within the hierarchical clusters, indicating that both methods are able to capture, sometime, the same underlying conformational states.

The inter-cluster graph (see Figure S6), where strong blue-violet edges connect most hierarchical clusters, demonstrating cluster’s high mutual similarity and confirming that they do not represent fundamentally different conformations.

Dimensionality reduction further supported this view: MDS preserved the global topology of the landscape, while UMAP and t-SNE highlighted local neighbourhoods, together revealing a compact, continuous conformational space rather than discrete states (see Figure S5). More precisely, the result similarity between two independent clustering methods demonstrate that the Ail-Vitronectin interface take a stable interface conformational with only subtle local fluctuations detectable by the finer hierarchical partition.

#### 3.1.c. Interactions at the Interface

To go beyond a simple analysis of the scores, we sought to determine whether the subtle differences revealed by the clustering methods were significant and not drive only by structural changes (RMSDs component of the equation). To do so, we analysed the noncovalent interaction (contact component in the equation) profiles of each group using MICLOT^38^. The interaction fingerprints obtained from DBSCAN and hierarchical clustering were remarkably consistent (Figure S7, A and B). both dominated by charge repulsions and hydrophobe-hydrophile repulsions, indicators of assembly quality showing the absence of unfavourable clash between residues. There is also a small contribution from salt bridges and arginine-aromatic contacts. The Van der Waals contact (Figure S7, C and D) counts similarly showed comparable trends across clusters, with overlapping error bars, confirming that the global interaction network remains stable regardless of the clustering.

However, closer inspection revealed few differences that a global analysis would overlook: the Arg374-Phe54 arginine–aromatic interaction was observed in only one specific cluster (see Figure 5, E and F), whereas Arg372-Phe54, which also involves Phe54, was conserved across all clusters. This indicates that MDQ can resolve local, transient contacts within an otherwise stable interface. These findings demonstrate that even in a globally stable system, the MDQ score is sensitive enough to distinguish between closely related clusters and to identify specific interaction fluctuations. Overall, these results show that MDQ not only validates interface stability at the global level but also captures small conformational and interactional variations along the trajectory, highlighting its precision for structural analysis.

### 3.2. Cytoplasmic domain of the type I TGF beta receptor / FKBP12 complex

#### 3.2.a. Overall stability and temporal dynamics

The TGFBR1-FKBP12 complex (Figure 6A) is a well know system, already used in the CAPRI-Q^13^ benchmark. The 2D MDQ heatmaps computed independently for each of the three replicates provide a direct visualization of interface global stability over time. In replicate 1, three discrete blocks are apparent along the diagonal, suggesting successive interface small reorganizations during the trajectory (Figure 6B). The similarity between the frames from 0 to 350 ns and 600-1000 ns shows that the transition at 300-600 ns is merely a transitional state. However, the score value for this transitional state remains between 0.4 and 0.6, indicating moderate similarity. The transitions remain of limited amplitude and do not correspond to major conformational switches. It is not visible to the other two replicants, indicating that the transition an artifact. Replicates 2 and 3 (Figure S8) each display only two weakly separated blocks, respectively around 600ns and 770 ns, with even lower contrast, further supporting the view that the interface undergoes only minor local fluctuations rather than global rearrangements. Overall, the consistently low MDQ values across all three replicates confirm that the TGFBR1-FKBP12 interface is globally stable and structurally converged, with only transient, low-amplitude rearrangements. Finally, the MDQ heatmaps resolve these subtle reorganizations, demonstrating that the metric is sensitive enough to capture fine dynamic events within an otherwise stable complex.

**Figure 6.**
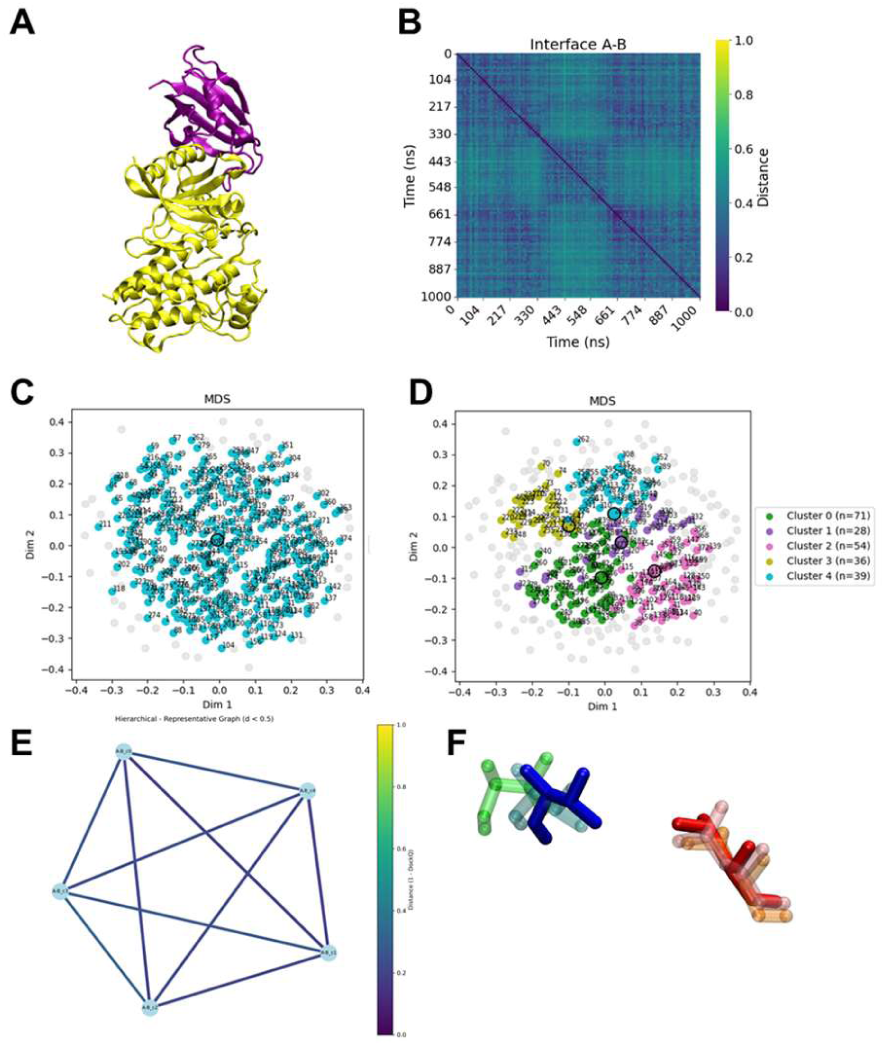
Comprehensive analysis of the TGFBR1-FKBP12 complex. **(A)** 3D structure of the complex (FKBP12 in purple, TGFBR1 in yellow). **(B)** 2D MDQ heatmap of the first replica trajectory (0 = identical, blue; 1 = distinct, yellow). **(C, D)** MDS projections of all frames colored by **(C)** DBSCAN clustering (1 cluster) and **(D)** hierarchical clustering (5 clusters). **(E)** Inter-cluster similarity graph for hierarchical clusters; edges are shown only for MDQ score < 0.5 and colored by score. **(F)** Leu276-Thr85 interfacial residue pair across representative hierarchical cluster structures.

#### 3.2.b. Conformational organization and clustering

To evaluate the conformational landscape of the TGFBR1-FKBP12 interface, we applied both DBSCAN and hierarchical clustering to the MDQ score matrix (Figure 6, C and D, and Figure S9). DBSCAN fail to identify more than one cluster in the merged trajectory. In contrast, hierarchical clustering partitioned the same data into five clusters, corresponding to fine subdivisions of a single stable interface state. MDS projection highlight continuous conformational space without clear separation between groups. The inter-cluster graph (Figure 6E) reinforced this interpretation: all nodes were interconnected by strong, dark-blue edges (MDQ < 0.5), confirming high mutual similarity among the hierarchical clusters.

#### 3.2.b. Conformational organization and clustering

Because DBSCAN identified only a single cluster, establishing an interaction profile was not possible for this method. Because the resulting curve of single cluster therefore can’t reflect distinct conformational states. That is why no conclusion can be drawn for DBSCAN. In contrast, results from the hierarchical clustering can be used to compare interaction patterns across the five sub-clusters. The interaction profiles (Figure S9) were remarkably consistent across all hierarchical clusters, while few variations can be seen. Van der Waals contact counts overlapping within error bars, confirming that the overall interaction network is conserved throughout the trajectory.

The Leu276-Thr85 residue pair (Figure 6F) was detected in all hierarchical sub-clusters. Although this pair adopted a similar spatial position across clusters, an actual interaction was observed only for one specific color-coded pair, indicating a transient, local contact superimposed on an otherwise stable interface. This demonstrates that MDQ, combined with hierarchical partitioning, is sensitive enough to resolve subtle interaction fluctuations even within a globally stable system where DBSCAN perceives only a single conformational state.

## 4. CONCLUSIONS AND PROSPECTS

This work introduces the MDQ score, a metric designed for comparing protein-protein interfaces in molecular dynamics simulations, building upon the DockQ framework traditionally used for docking model evaluation. Unlike the original score, the MDQ score was redefined to ensure symmetry, producing identical results regardless of the order of compared structures (*i,j* or *j,i*). By incorporating residue contact pairs into the scoring scheme, the MDQ score captures non-bonding interactions within interfaces, offering a more precise and nuanced alternative to conventional C_α_-RMSD analyses and clustering. Its sensitivity allows the detection of subtle conformational and contacts changes in stable interfaces, making it a powerful tool for interface analysis.

As demonstrated here, the score proves its precision and utility for analyzing complete interfaces. However, its applications extend beyond this: it could also be applied to distinct interface regions (*core*, *support*, *rim*)^41^, enabling the assessment of each region’s role in interface stability, as previously shown^38^.

However, further refinements are possible. Mathematically, explore alternative loss function for RMSD correction could improve accuracy, particularly for low-overlap cases. Additionally, integrating side-chain iRMSD^13^ (beyond backbone atoms) and other MD-derived descriptors: such as Buried Surface Area (BSA) or binding free energy variations, could enhance the metric’s discriminative power.

## Software availability

The script is implemented in Python 3 and relies on MDTraj^42^, enabling compatibility with all molecular dynamics trajectory file formats supported by the library. It is integrated into MICLOT^38^, a software developed in previous research for protein molecular interaction analysis. MDQ script is open-source and freely available with its documentation on GitHub: https://github.com/TMiclot/MICLOT

## ASSOCIATED CONTENT

### Supporting Information

The following files are available free of charge.

The Supporting Information (PDF file) provides additional methodological details and extended validation data.

## AUTHOR INFORMATION

### Author Contributions

The manuscript was written through contributions of all authors. All authors have given approval to the final version of the manuscript.

Jenny Ehles: Software development; MD simulation; formal analysis; data curation.

Florent Barbault: MD simulation; writing – review and editing; supervision.

Tom Miclot: Conceptualization; writing – review and editing; supervision.

### Funding Sources

Any funds used to support the research of the manuscript should be placed here (per journal style).

### Notes

The authors declare no conflict of interest.

## ACKNOWLEDGMENT

CNRS (Centre National de la Recherche Scientifique) is gratefully acknowledged for his financial support of this work through *Dotation initiale de démarrage*. This project was provided with computer and storage resources by GENCI (Grand Equipement National de Calcul Intensif) at IDRIS thanks to the grant 2026-AD010716281R1 on the supercomputer Jean Zay’s V100 and CSL partition. We acknowledge the use of the LLM (large language model) CNRS Emmy - Mistral AI to improve the grammatical and stylistic accuracy of the manuscript.

## Supporting Information

## I. DockQ score is not symmetric

The DockQ score is not symmetric when swapping the model and native inputs, as demonstrated by the differing results obtained from the command `DockQ <model> <native>` versus `DockQ <native> <model>`. This asymmetry arises from the internal calculations of iRMSD, LRMSD, fnat,and fnonnat which depend on the alignment direction.

This phenomenon can be demonstrated by conducting tests on structures based on examples from GitHub (https://github.com/wallnerlab/DockQ). In Table S1, swapping the model and native structures yields DockQ scores of 0.941 and 0.954, respectively, despite identical LRMSD values. Similarly, Table S2 shows significant variations in DockQ, iRMSD, and fnat when the model and native are interchanged, further illustrating this directional dependency.

**Table S1.** DockQ scores computed for two PDB structures (2HHB_hem.cif and 1HHO_hem.cif) with model and native inputs swapped. The asymmetry in DockQ, iRMSD, and fnat is evident despite identical LRMSD values.

| Model | **2HHB_hem.cif** | **1HHO_hem.cif** |
|---|---|---|
| Native | **1HHO_hem.cif** | **2HHB_hem.cif** |
| **Total DockQ** | **0.941** | **0.954** |
| Native chains | A, B | C, D |
| Model chains | C, D | A, B |
| DockQ | 0.941 | 0.954 |
| iRMSD | 0.486 | 0.420 |
| LRMSD | 1.481 | 1.481 |
| fnat | 0.947 | 0.964 |
| fnonnat | 0.036 | 0.053 |
| F1 | 0.956 | 0.956 |
| clashes | 0 | 0 |

**Table S2.** DockQ scores for the 1A2K complex with different chain mappings between model and native structures. The results highlight variations in DockQ, iRMSD, LRMSD, and fnat when the model and native inputs are reversed, demonstrating the score’s lack of symmetry.

| Model | **1A2K_r_l_b.model.pdb** | **1A2K_r_l_b.pdb** |
|---|---|---|
| Native | **1A2K_r_l_b.pdb** | **1A2K_r_l_b.model.pdb** |
| **Total DockQ** | **0.653** | **0.798** |
| Native chains | A, B | A, B |
| Model chains | B, A | B, A |
| DockQ | 0.994 | 0.997 |
| iRMSD | 0.000 | 0.000 |
| LRMSD | 0.000 | 0.000 |
| fnat | 0.983 | 0.992 |
| fnonnat | 0.008 | 0.017 |
| F1 | 0.987 | 0.987 |
| clashes | 0 | 0 |
| Native chains | A, C | A, C |
| Model chains | B, C | B, C |
| DockQ | 0.511 | 0.623 |
| iRMSD | 1.237 | 1.646 |
| LRMSD | 6.864 | 8.131 |
| fnat | 0.333 | 0.893 |
| fnonnat | 0.000 | 0.500 |
| F1 | 0.500 | 0.641 |
| clashes | 0 | 0 |
| Native chains | B, C | B, C |
| Model chains | A, C | A, C |
| DockQ | 0.453 | 0.775 |
| iRMSD | 2.104 | 0.938 |
| LRMSD | 8.131 | 6.864 |
| fnat | 0.500 | 1.000 |
| fnonnat | 0.107 | 0.667 |
| F1 | 0.641 | 0.500 |
| clashes | 0 | 0 |

## II. Discussion on the choice of the mathematical form of the loss function

### II.1. Overview of the *loss* function

The logarithmic loss function, below, is particularly well-suited for this application as it directly addresses the non-linear relationship between atomic overlap and RMSD reliability. It is tailored to structural biology comparisons involving continuous similarity metrics. The function penalizes low-overlap comparisons—where RMSD values based on few shared atoms are inherently unreliable—by reaching its maximum value of approximately 50.83 when no atoms are shared (*frac(i,j)* = 0). Conversely, it approaches its minimum value of 1 when *frac(i,j)* ≈ 1, ensuring that RMSD remains largely uncorrected when atomic overlap is high. See equation below:

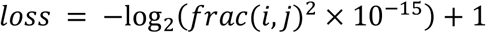

The logarithmic form enhances sensitivity to small changes in *frac(i,j)* at low values while being less responsive at high values, aligning with the intuition that minimal atomic alignment requires stronger correction. Additionally, the function is continuous and differentiable over the interval [0, 1], avoiding abrupt jumps, with the small constant 10^−1^^5^ preventing numerical instability at *frac(i,j)* = 0. See Figure S1.

**Figure S1.**
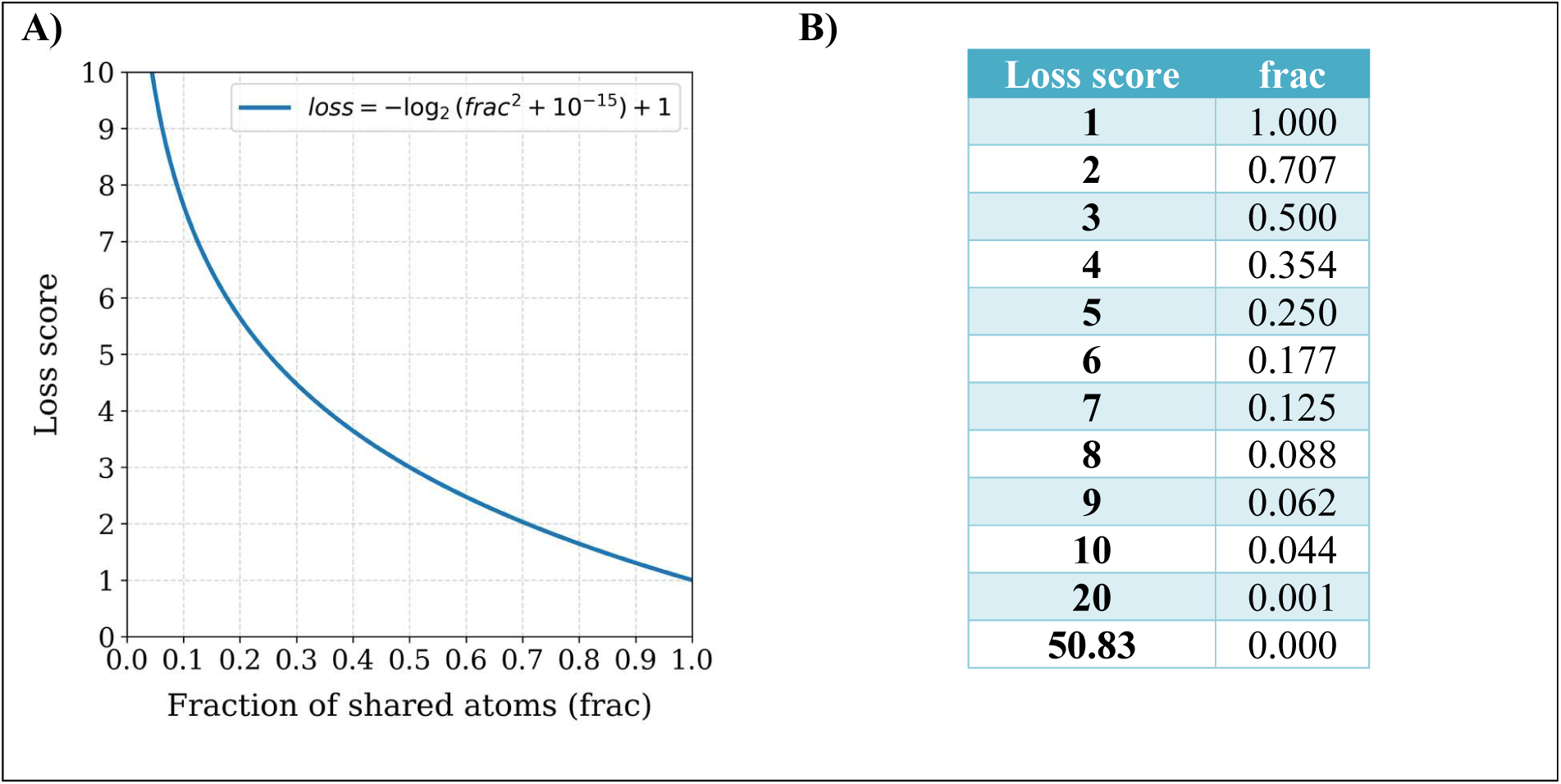
For clarity, frac(i,j) is simplified to frac throughout the discussion. **A)** The loss function exhibits a logarithmic decay, decreasing rapidly at low frac values with a steep initial slope, then gradually flattening as frac approaches 1. **B)** This behaviour is summarized in the accompanying table, which reports representative loss values for corresponding frac values, including the two extrema: frac = 1 (loss = 1) and frac = 0 (loss ≈ 50.83).

### II.2. Critics of the *loss* function

Two primary criticisms can be raised regarding the loss function’s design: the choice of the logarithmic base and the arbitrary offset. The offset of 10^−15^ serves as a numerical safeguard to prevent division by zero but lacks a clear physical or statistical justification. A smaller or larger value could subtly alter the function’s behavior for small values of frac(*i,j*), potentially affecting the correction’s sensitivity. Additionally, the use of base-2 logarithm is arbitrary; while alternative bases would preserve the qualitative behavior, they would scale the magnitude of the correction differently. The choice of base thus primarily influences the intensity of the penalty rather than its fundamental behavior.

### II.3. Discussion on alternative functions

Other methods could be considered, but none are inherently superior for this specific problem. A reciprocal function would similarly penalize low frac values, but it decays more aggressively than the logarithmic form, potentially overcorrecting RMSD for moderately low overlap. Exponential functions could also be used, but they lack the intuitive scaling of the logarithmic approach and may require tuning of terms parameters. See Figure S3.

The current logarithmic loss function is well-justified because it naturally aligns with information-theoretic principles and provides a smooth, interpretable correction that avoids extreme values while strongly penalizing unreliable comparisons. Unless there is a specific reason to favor a different equation type, the logarithmic approach remains a strong and theoretically sound choice. However, it is possible to identify the most appropriate type of function and parameterise its terms by using a large database of molecular dynamics trajectories for various biological systems. But this last is out of the scope of the current work.

**Figure S3.**
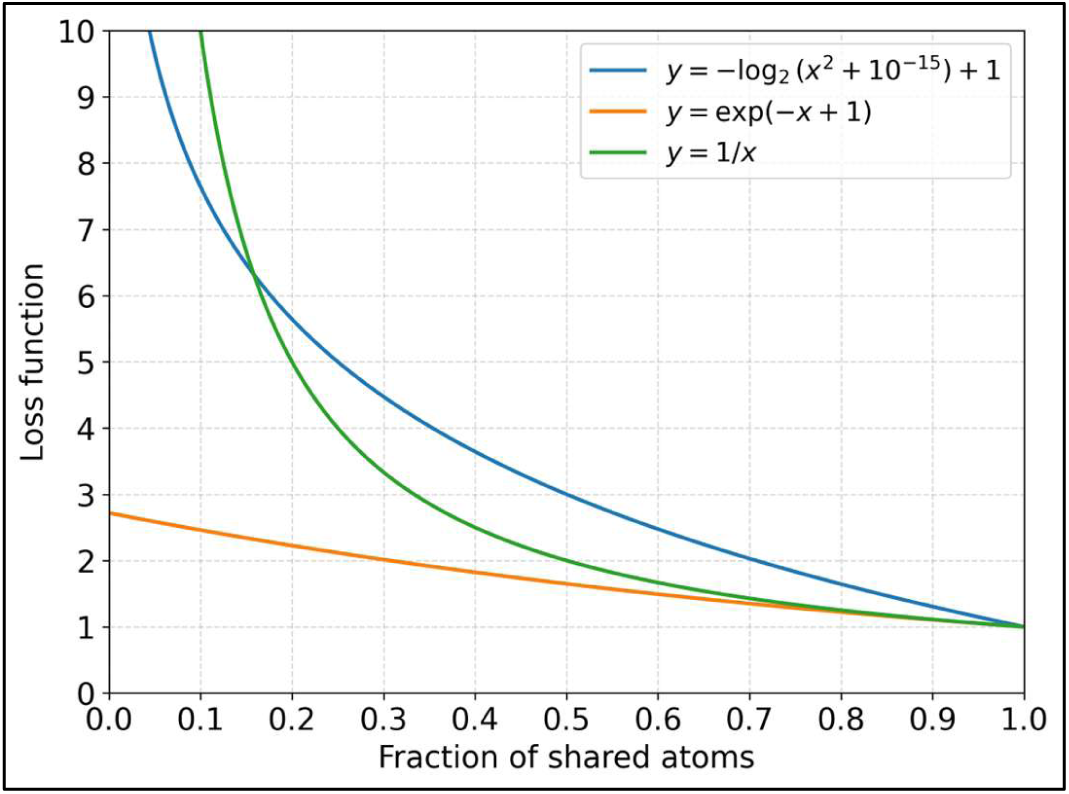
The figure displays three curves: the selected loss function and two potential analogous functions (reciprocal and exponential). All three functions converge to a value of 1 at x = 1, as intended. The reciprocal function initially lies below the loss function but intersects it at x ≈ 0.158 (y ≈ 6.320), after which it rises above the loss curve. In contrast, the exponential function remains consistently below the loss function across the entire range. A simple parameterization was applied to ensure the exponential curve passes through y = 1 at x = 1.

## III. Critics of the rescaling method for low RMSD values

### III.1. Limitation of the multiplicative correction

The current formulation, below, applies a pure multiplicative correction, where the RMSD is scaled proportionally to the loss function. When frac(*i,j*) = 1, this mean that RMSD_w_ is exactly equal to RMSD, preserving the original value:

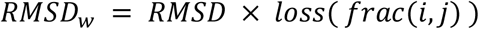

However, the rescaling method for RMSD presents notable limitations, particularly for low RMSD values. More precisely, for low frac(*i,j*) value, small RMSD values remain low after the rescaling and potentially misrepresent good structural alignment. This highlights the need for a dedicated treatment of low RMSD values, which would require extensive analysis of multiple trajectories across diverse systems to define appropriate ranges.

***Example:*** *If an RMSD has a excellent value of 0.1 Å but the fraction of shared atoms frac(i,j) is only 0.125 (i.e., ∼12.5% of atoms are shared and used for RMSD computation), the rescaled RMSD_w_ becomes: 0.1 × 7 ≈ 0.7 Å, which falsely suggests a good result*.

### III.2. Another possible correction equation

Introducing additive parameters can mitigate this issue and an alternative equation can be:

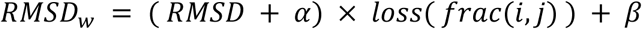

The additive terms α and β provide greater control over the rescaling behaviour: α shifts the RMSD before scaling, while β adjusts the baseline. This can prevent insufficient inflation of low RMSD values when frac is small, as demonstrated in the Figure S4.

This additive rescaling function effectively addresses low RMSD values but disproportionately inflates high RMSD values. To mitigate this, a hybrid approach can be implemented by combining both multiplicative and additive formulations through a threshold ω. Specifically, the multiplicative form is retained for ω < RMSD, while the additive form is applied for a value in the range: 0 < RMSD < ω.

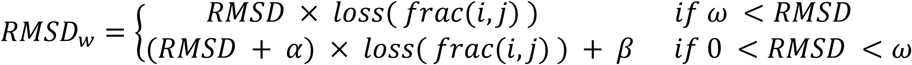

This hybrid rescaling ensures that low RMSD values are corrected without excessive amplification, while high RMSD values remain proportionally scaled, thus balancing accuracy across the full range of structural deviations.

However, the parametrisation of additive terms α and β would require systematic benchmarking on large molecular dynamic dataset. This is not in the scope of the current work.

**Figure S4.**
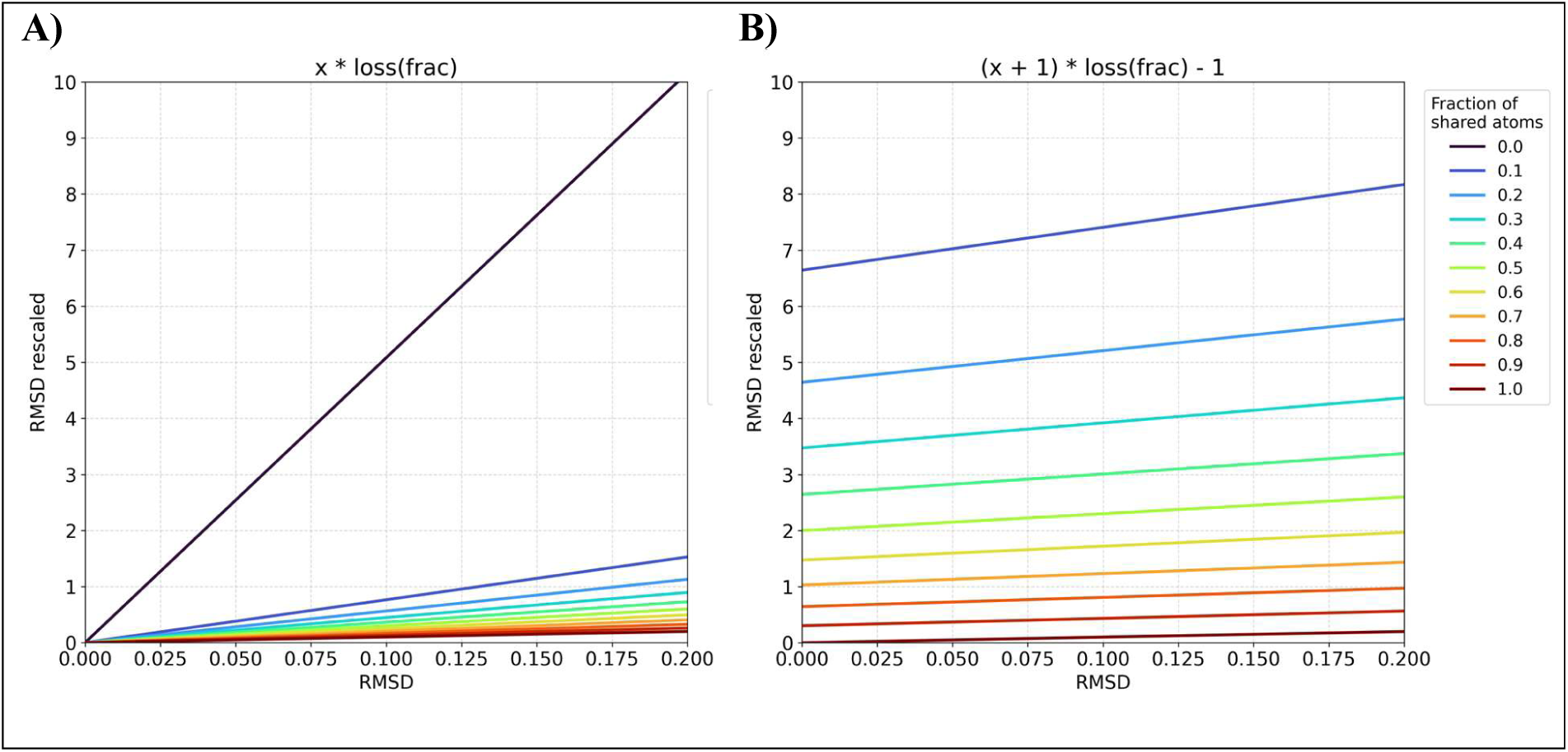
The figure illustrates how RMSD values are corrected to RMSD_w_ using the described functions, with multiple curves plotted for varying values of frac (i.e. frac(*i,j*)) ranging from 0 to 1 in increments of 0.1. The x-axis range for RMSD was arbitrarily set to [0, 0.2] to focus on low RMSD values. Panel **A)** displays the curves for the multiplicative equation, where all lines pass through zero. The curve for frac = 0 imposes the strongest penalty on low RMSD values, while the other curves show minimal correction, even for low frac values. Panel **B)** presents the curves for the additive equation, where no line passes through zero, and the curve for frac = 0 falls outside the graph. In this case, increasingly severe penalties are applied for low frac values, resulting in RMSD_w_ values exceeding 7.

### III.3. Properties and parametrisation of correction equations - Generality

While alternative correction functions could be identified, their development would require rigorous analysis of extensive molecular dynamics trajectory datasets across diverse biological systems. Beyond identifying an optimal functional form, the process would also demand careful parameterization of its terms. Furthermore, any proposed correction function and its terms must remain compatible with the existing d_1_ = 8.5 Å for LRMS and d_2_ = 1.5 Å for iRMS parameters from the DockQ score^8,9^; otherwise, additional reparameterization efforts would be necessary to maintain consistency for d_1_ and d_2_. Using a multiplicative correction maintains a proportional relationship with the initial value, limiting the impact and preserving the original d_1_ and d_2_ parameter values.

Beyond identifying the optimal correction equation, a significant effort is required to determine the best value for the threshold parameter ω, which discriminates between high and low RMSD values.

### IV. Variants of the MDQ Score Equation

The MDQ score could be further refined by integrating additional structural and energetic descriptors. The conservation of interface size, quantified through the fractional Buried Surface Area (FBSA) derived from BSA, represents a readily available metric in docking predictions that could have been incorporated into the original DockQ framework, but is not. Similarly, the stability of binding free energy, captured by FΔG and computable from molecular dynamics trajectories, offers another dimension for evaluation. Including these parameters would enhance the score’s robustness by accounting for both geometric consistency and thermodynamic stability, thereby providing a more holistic assessment of structural similarity. However, this enhancement is not without limitations, as it introduces dependencies on the specific methods used for SASA or free energy calculations, which can vary significantly across implementations, thereby affecting the robustness and reproducibility of the score.

### IV.1. Incorporating Buried Surface Area (BSA)

The Buried Surface Area (BSA), defined as the difference between the Solvent-Accessible Surface Area (SASA) of the isolated components and that of the complex, serves as a valuable metric for assessing protein complex stability. The size of an interface is already known to correlate with complex stability ^43,44^. Within the framework of the MDQ score, BSA can refine the evaluation by providing a nuanced metric for comparing protein complexes in molecular dynamics simulations, particularly in how the interface size changes between two conformations along the trajectory. The comparison between interfaces can be incorporated into the score using the fractional BSA (FBSA), which is defined as the ratio of the minimum to the maximum BSA values between two compared interfaces. This metric quantifies interface size proportionality, with values approaching 1 for structurally consistent interfaces and 0 for divergent ones. The use of FBSA allows the score to be applied not only to the entire interface but also to specific structural regions ^41^ (e.g., rim, core, or support), enhancing its adaptability.

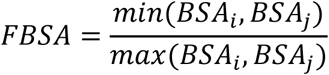

*MDQ Score equation incorporating the FBSA value:*

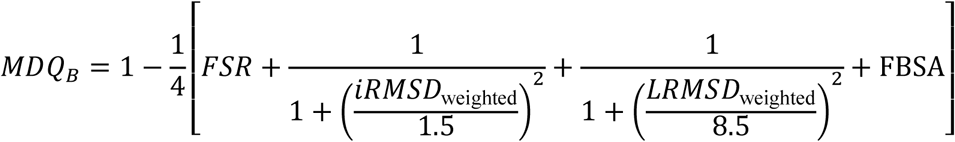

However, multiple methods exist for computing SASA, each introducing variability. Differences arise from the treatment of hydrogens—whether they are explicitly modelled, implicitly included, or omitted, as in the NACCESS algorithm^45^. Additionally, the choice of atomic van der Waals radii can influence the final value, even if the variation is small. For instance, the oxygen atom radius is reported as 1.40 Å by Chothia^46^ and 1.52 Å by Bondi^47^. Further discrepancies stem from the algorithm employed, whether it relies on mathematical tessellation^48^, Voronoi tessellation^49^, or statistical methods using linear empirical scaling factors^50^.

Given the multitude of SASA calculation methods, it would be advisable to test several to assess their respective effects. Additionally, parameterizing a constant value for FBSA—similar to what has been done for RMSD values in the original DockQ score^8,9^ —may be necessary. But such an investigation falls outside the scope of this research.

### IV.2. Incorporating Gibbs free energy (ΔG)

The binding affinity (ΔG) of a protein complex serves as a critical metric for assessing interaction strength, with more negative ΔG values indicating stronger binding. Within the framework of the MDQ score, ΔG can refine the evaluation by providing a nuanced metric for comparing protein complexes in molecular dynamics simulations, particularly in how binding affinity varies between two conformations along the trajectory. The comparison between complexes can be incorporated into the score using the fractional ΔG (FΔG), defined as:

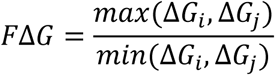

*MDQ Score equation incorporating the F*Δ*G value:*

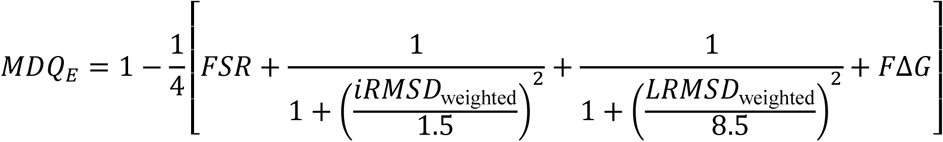

This metric quantifies the proportionality of binding affinities, with values approaching 1 for complexes with similar affinities and increasing as the affinities diverge. The use of FΔG allows the score to be applied to the overall binding affinity, enhancing its adaptability in comparative analyses. However, multiple methods exist for estimating binding free energy, each introducing variability. Differences arise from the underlying computational approach—whether based on end-point methods like MMPB(GB)SA^51^, contact-based predictions^52,53^. Additionally, in some methods the choice of force field parameters can influence the final free energy values, even if the variations are subtle. For instance, discrepancies may emerge from the treatment of solvation models or the definition of interaction cutoffs.

Given the multitude of free energy calculation methods, it would be advisable to test several to assess their respective effects. Additionally, parameterizing a constant value for free energy comparisons—similar to what has been done for RMSD values in the original DockQ score—may be necessary. However, such an investigation falls outside the scope of this research.

### IV.3. Incorporating both ΔG and BSA

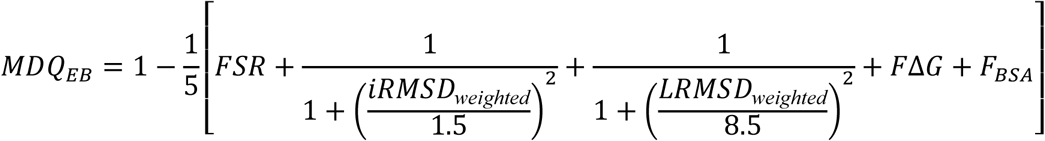

## V. Ail/vitronectin complex

## V.1. Cluster identification and representation

**Figure S5.**
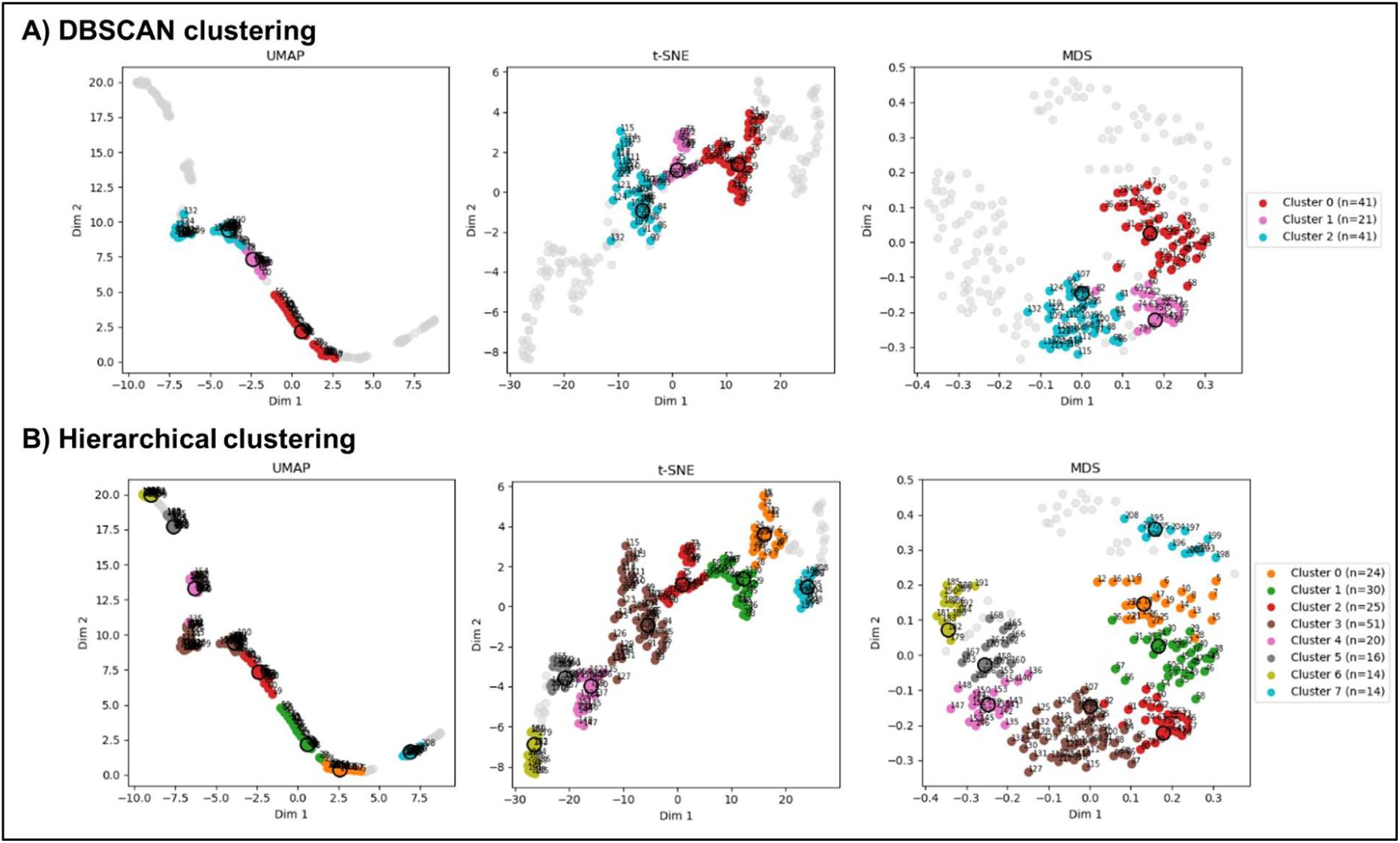
Dimensionality-reduction projections of the of the 2D MDQ analysis of Ail-Vitronectin trajectory colored by clustering assignment. **(A)** DBSCAN clustering (3 clusters) and **(B)** hierarchical clustering (8 clusters) visualized with UMAP (left), t-SNE (center), and MDS (right). Colored points represent frames assigned to each cluster; grey points denote noise/unassigned frames. Black circles indicate the representative structure of each cluster (i.e., the frame with the lowest mean MDQ distance to all other members).

**Figure S6.**
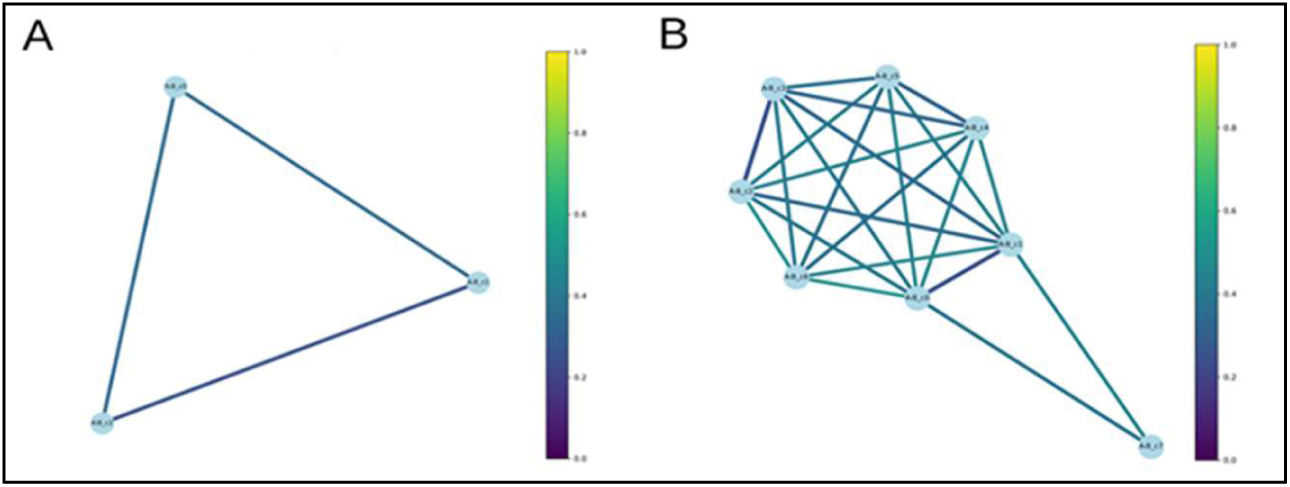
Inter-cluster similarity graphs for the Ail-Vitronectin complex. Each node represents the representative structure of a cluster (the frame with the lowest mean MDQ distance to all other members). (A) DBSCAN clustering (3 clusters); (B) hierarchical clustering (8 clusters). Edges connect cluster representatives with an MDQ distance < 0.5 and are colored according to the pairwise MDQ score (blue = highly similar, yellow = distinct).

## V.2. Non-bonding interaction stabilising the stability of the complex

**Figure S7.**
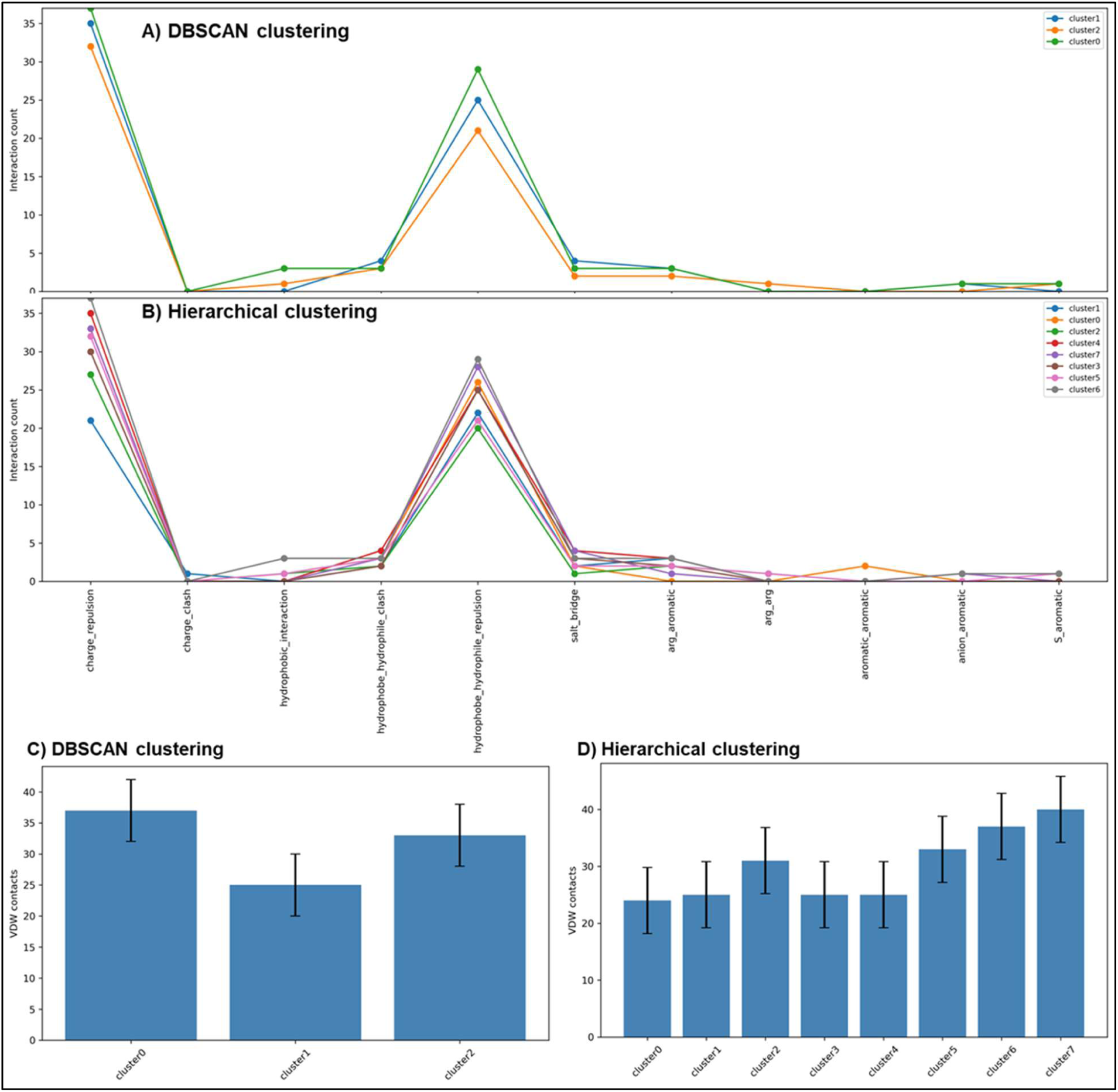
Non-covalent interaction analysis of the Ail–vitronectin interface across clusters. **(A, B)** Interaction profiles showing the count of each interaction type (charge repulsion/clash, hydrophobic, hydrophobe–hydrophile clash/repulsion, salt bridges, arginine–aromatic, arginine–arginine, aromatic–aromatic, anion–aromatic, sulfur–aromatic) for each cluster identified by **(A)** DBSCAN (3 clusters) and **(B)** hierarchical clustering (8 clusters). **(C, D)** Mean number of Van der Waals contacts per cluster, all structure in the same cluster where analysed, (± standard deviation) for **(C)** DBSCAN and **(D)** hierarchical clustering.

## VI. Cytoplasmic domain of the type I TGF beta receptor / FKBP12 complex

## V.1. Evaluation of trajectory convergence

**Figure S8.**
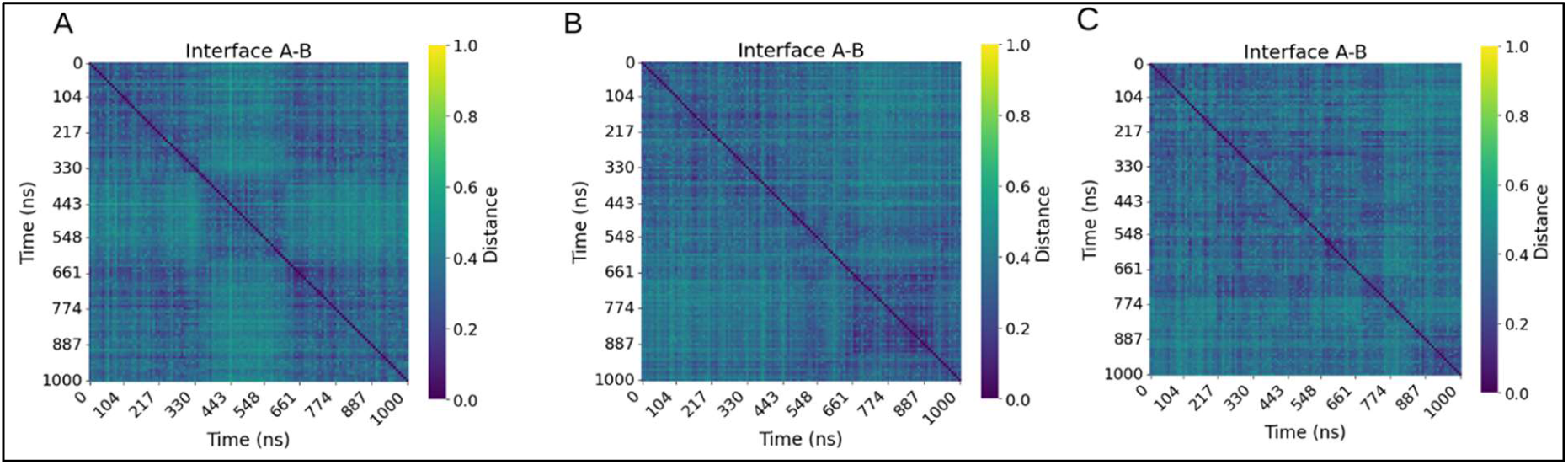
2D MDQ heatmaps of the three independent replicates of the TGFBR1-FKBP12complex. MDQ score heatmap computed between all frames of **(A)** replicate 1, **(B)** replicate 2, and **(C)** replicate 3, with time (ns) on both axes. The color scale ranges from 0 (identical interfaces, dark blue) to 1 (distinct interfaces, yellow). Blocks along the diagonal indicate periods of conformationally similar interface states. Replicate 1 displays three blocks, but replicates 2 and 3 show only weakly separated blocs, consistent with a globally stable interface.

## V.2. Cluster identification and representation

**Figure S9.**
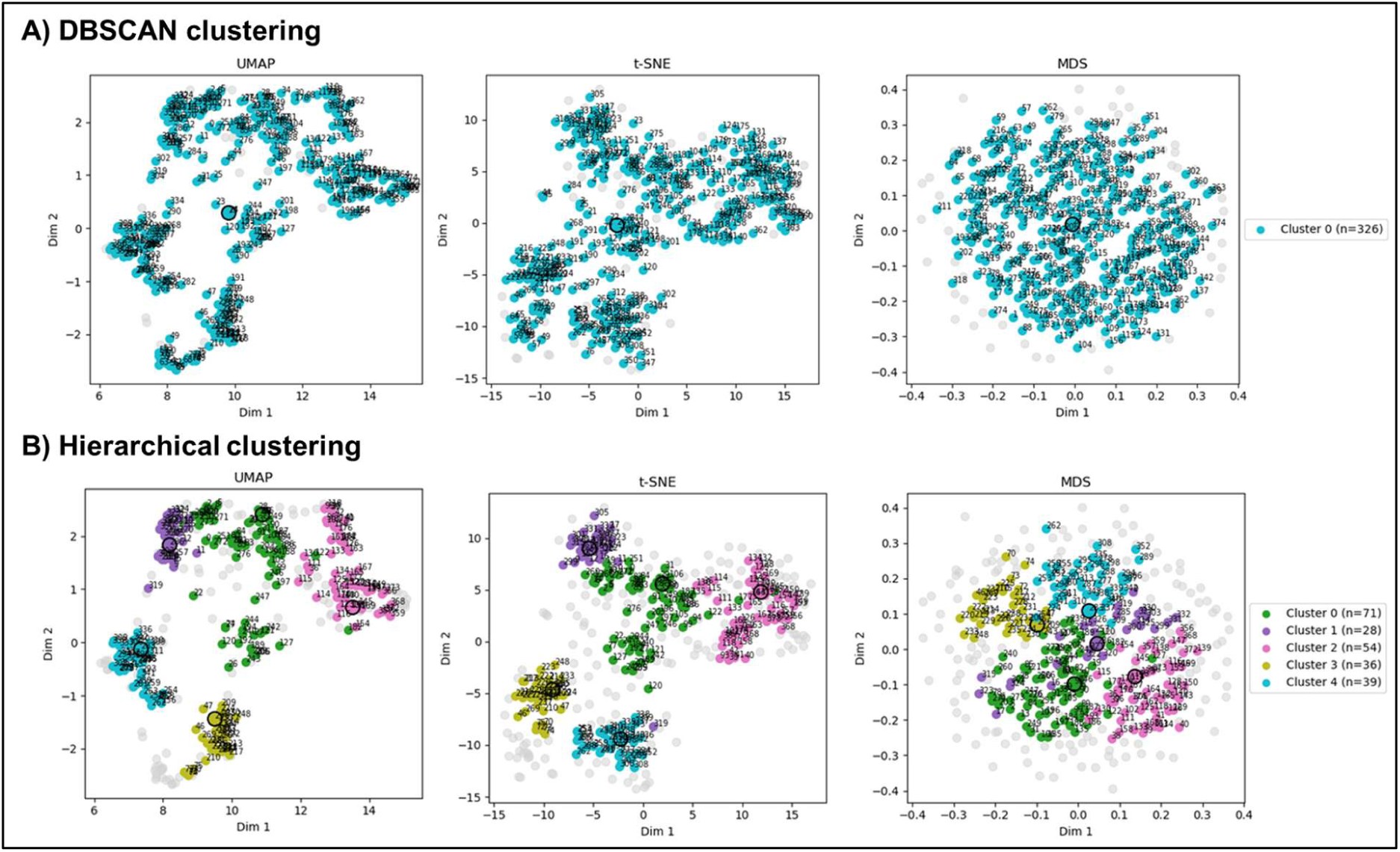
Dimensionality-reduction projections of the 2D MDQ analysis of TGF beta receptor / FKBP12 complex trajectory colored by clustering assignment. **(A)** DBSCAN clustering (1 clusters) and **(B)** hierarchical clustering (5 clusters) visualized with UMAP (left), t-SNE (center), and MDS (right). Colored points represent frames assigned to each cluster; grey points denote noise/unassigned frames. Black circles indicate the representative structure of each cluster (i.e., the frame with the lowest mean MDQ distance to all other members).

## V.3. Non-bonding interaction stabilising the stability of the complex

**Figure S10.**
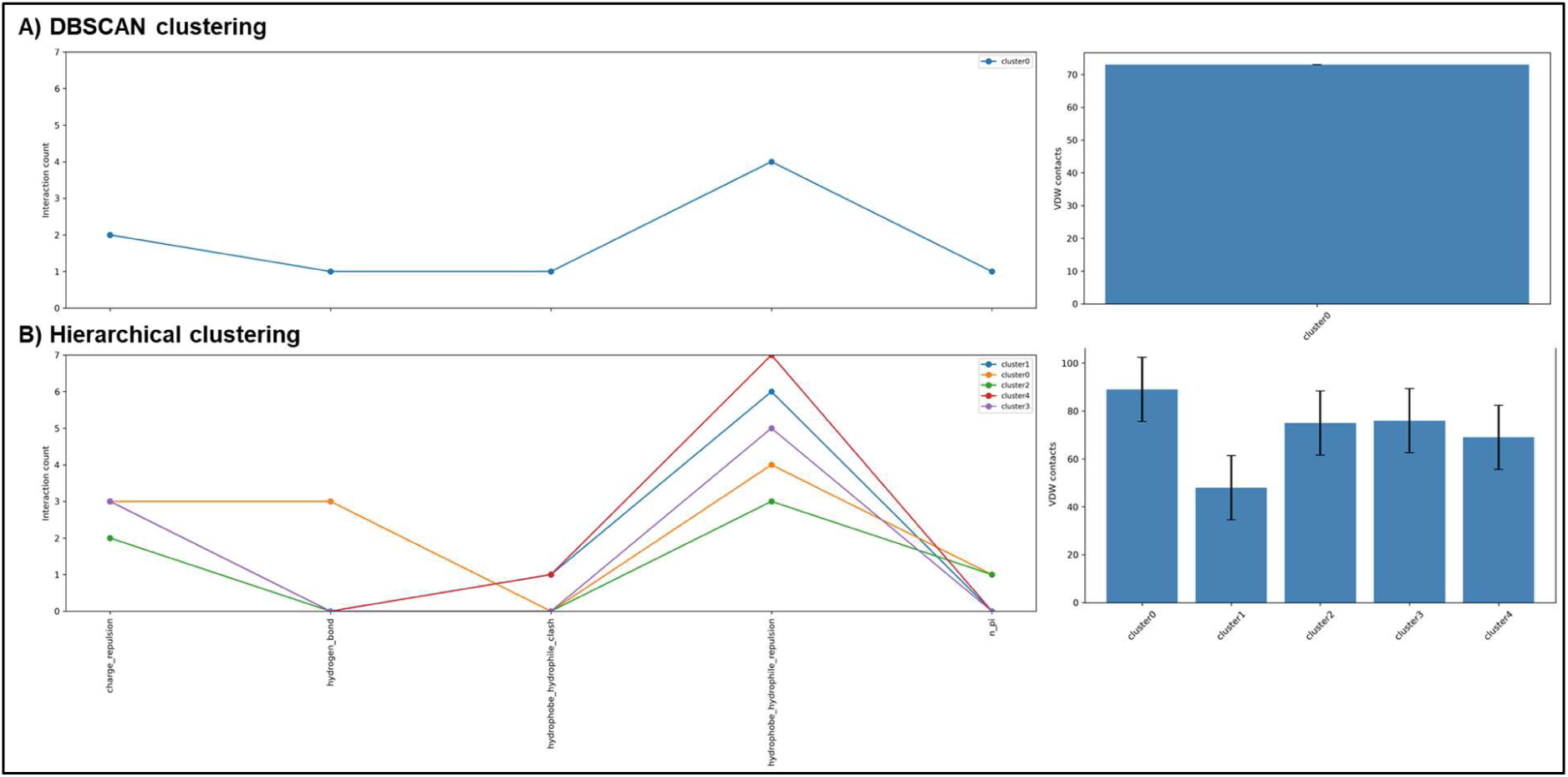
Non-covalent interaction analysis of the FKBP12-TGFBR1 interface across clusters. **(A)** DBSCAN clustering (single dominant cluster): interaction count profile by type (left) and Van der Waals contact count (right). **(B)** Hierarchical clustering (5 sub-clusters): per-cluster interaction profiles (left) and mean VdW contacts ± SD (right).

## Notes

### Competing Interest Statement

The authors have declared no competing interest.

